# When less is more – A fast TurboID KI approach for high sensitivity endogenous interactome mapping

**DOI:** 10.1101/2021.11.19.469212

**Authors:** Alexander Stockhammer, Carissa Spalt, Laila S. Benz, Shelly Harel, Vini Natalia, Lukas Wiench, Christian Freund, Benno Kuropka, Francesca Bottanelli

## Abstract

In recent years, proximity labelling has established itself as an unbiased and powerful approach to map the interactome of specific proteins. While physiological expression of the labelling enzyme is beneficial for the mapping of interactors, generation of the desired cell lines remains time-consuming and challenging. Using our established pipeline for the rapid generation of C- and N-terminal CRISPR-Cas9 knock-ins (KIs) based on antibiotic selection, we were able to compare the performance of commonly used labelling enzymes when endogenously expressed. Endogenous tagging of the μ subunit of the AP-1 complex with TurboID allowed identification of known interactors and cargo proteins that simple overexpression of a labelling enzyme fusion protein could not reveal. We used the KI-strategy to compare the interactome of the different adaptor protein (AP) complexes and clathrin and were able to assemble lists of potential interactors and cargo proteins that are specific for each sorting pathway. Our approach greatly simplifies the execution of proximity labelling experiments for proteins in their native cellular environment and allows going from CRISPR transfection to mass spectrometry analysis and interactome data in just over a month.

## Introduction

The biological role of a protein is shaped by its subcellular localization and its interaction with other biomolecules. Therefore, mapping the interactors of a given protein can be crucial for understanding its biological function. In the past several years, proximity labelling with biotin in living cells has emerged as a complementary approach to classic affinity purification/mass spectrometry (AP/MS)-based methods for mapping protein-protein interactions in living cells and organisms^1,2^. The proximity labelling is carried out by enzymes genetically fused to the protein of interest (POI) that catalyze the formation of a highly reactive biotin intermediate labelling proteins within a small radius (1-10 nm)^3,4^ in a promiscuous manner. A key advantage of proximity labelling-based interactome mapping compared to traditional approaches is that very weak and transient interactions can also be captured. The biotinylation itself provides a unique chemical moiety that can be used for subsequent enrichment and identification.

The enzymes used for proximity labelling are either engineered peroxidases (APEX^5^, APEX2^6^) or engineered biotin ligases (BioID^7^, BioID2^8^, TurboID^9^ and miniTurboID^9^). APEX and APEX2 use H_2_O_2_ as a co-substrate to rapidly generate a highly reactive phenoxyl radical from biotin-phenol that reacts specifically with electron-rich side chains (primarily tyrosine)^9^. An attractive feature of APEX peroxidases is the fast labelling kinetics (labelling time: <1 min) that enable probing with a high temporal resolution. However, on the downside, they require H_2_O_2_, which causes oxidative stress in living cells and thus cannot be used for proximity labelling in living organisms. In contrast, biotin ligases simply need non-toxic, highly soluble biotin as a substrate which in an ATP-dependent reaction, is converted into a reactive biotinoyl-5’-AMP intermediate that covalently tags proximal lysine residues^7^. Although the labelling time could be reduced from >18h for BioID^7^ to less than an hour with TurboID^9^, labelling with biotin ligases is significantly slower than with APEX peroxidases.

To avoid artifact as a result of overexpression, such as mislocalization and protein aggregation among others^10-12^, physiological expression levels of the POI fusion protein are preferred for interactome mapping experiments. Moreover, less abundant interactors that would be hidden in the unspecific labelling background that is caused by non-physiological expression levels might be identified this way. To control expression levels of the fusion protein overall, various different approaches have been implemented. These include the use of small-molecule induced expression systems in a safe DNA locus (such as the Flp-In™ T-Rex™ system)^13,14^, CRISPR-Cas9 based genome editing^15,16^ or knock out and replacement approaches in which the gene is first knocked out and then the fusion protein is integrated in a safe locus^17^. However, these methods either require the time-consuming selection and testing of single cell clones (CRISPR-based approaches) or are limited to standardized commercially available cell lines and not endogenous protein expression (inducible expression systems).

Here, we combine a rapid KI strategy for C- or N-terminal tagging based on antibiotic selection of positively edited cells^18,19^ with proximity-based proteomics to detect interactors at physiological expression levels. This versatile approach enabled us to endogenously tag proteins with the four most commonly used labelling enzymes (APEX2, BioID2, miniTurboID and TurboID) and compare their performance when expressed endogenously. In the past, proximity biotinylation approaches have been instrumental to map the interactome of membrane-bound organelles and membrane associated proteins^20-24^. Therefore, we tagged the μ1A subunit of the adaptor protein complex AP-1 (AP1μA) at the C-terminus and clathrin light chain (CLC) at the N-terminus. AP-1, together with CLC, mediates specific transport between the *trans*-Golgi network (TGN) and endosomes^25,26^ and transiently associates to membranes, leaving excess, non-membrane bound AP-1 subunits and CLC in the cytosol. Our data reveals that endogenous tagging allows proximity labelling with higher specificity compared to simple overexpression of the labelling enzyme. Known interactors of AP-1 as well as specific cargo were more highly enriched or even exclusively found in the experiments performed with a KI cell line. We identified TurboID to be best suited for knock-in proximity labelling and propose a pipeline for rapid endogenous tagging to improve the workflow and quality of proximity-based mass spectrometry experiments. The use of this pipeline allowed us to compare the interactome of different AP complexes, including the low-abundant AP-4 and assemble a list of potential interactors and cargo proteins for each adaptor protein complex. Lastly, we compared the interactome of CLCa to AP-1 and AP-2, which are both clathrin adaptors, to show that the here presented KI-strategy with TurboID allows identification of pathway specific interactors.

## Results

### Endogenous C-terminal tagging of AP1μA with labelling enzymes

To evaluate the performance of commonly used labelling enzymes, we used the CRISPR-Cas9 system to genetically fuse APEX2, BioID2, miniTurboID and TurboID to the C-terminus of our chosen target AP1μA. The insertion of a geneticin (G418) resistance cassette downstream of a polyadenylation signal into the targeted gene locus (AP1M1) allowed us to rapidly select for cells that were successfully modified^19^. In addition, a V5-tag was inserted for KI validation via western blot and immunofluorescence. The CRISPR strategy is illustrated in Fig. 1A. Cells were transfected with plasmids encoding for a gRNA targeting the genomic locus and homology repair plasmids containing sequences of the various labelling enzymes. Three days post transfection, G418 was added, and cells were allowed to grow back to confluency to perform downstream analysis. Expression of the fusion proteins in the four KIs was validated with western blot and immunofluorescence (Fig. 1B, C). Homology-directed repair is a rare event, mainly yielding a mixed population of heterozygous cells. Therefore, we manually evaluated the overall efficacy of each KI using immunofluorescence microscopy to detect positively edited cells and found KI rates of 43% (APEX2), 63% (BioID2), 45% (MiniTurboID), 59% (TurboID). After comparison to the expression levels of edited AP1μA on the western blot, we assume that most of the cells in our population are heterozygous for AP1μA fusion protein expression.

**Fig. 1:**
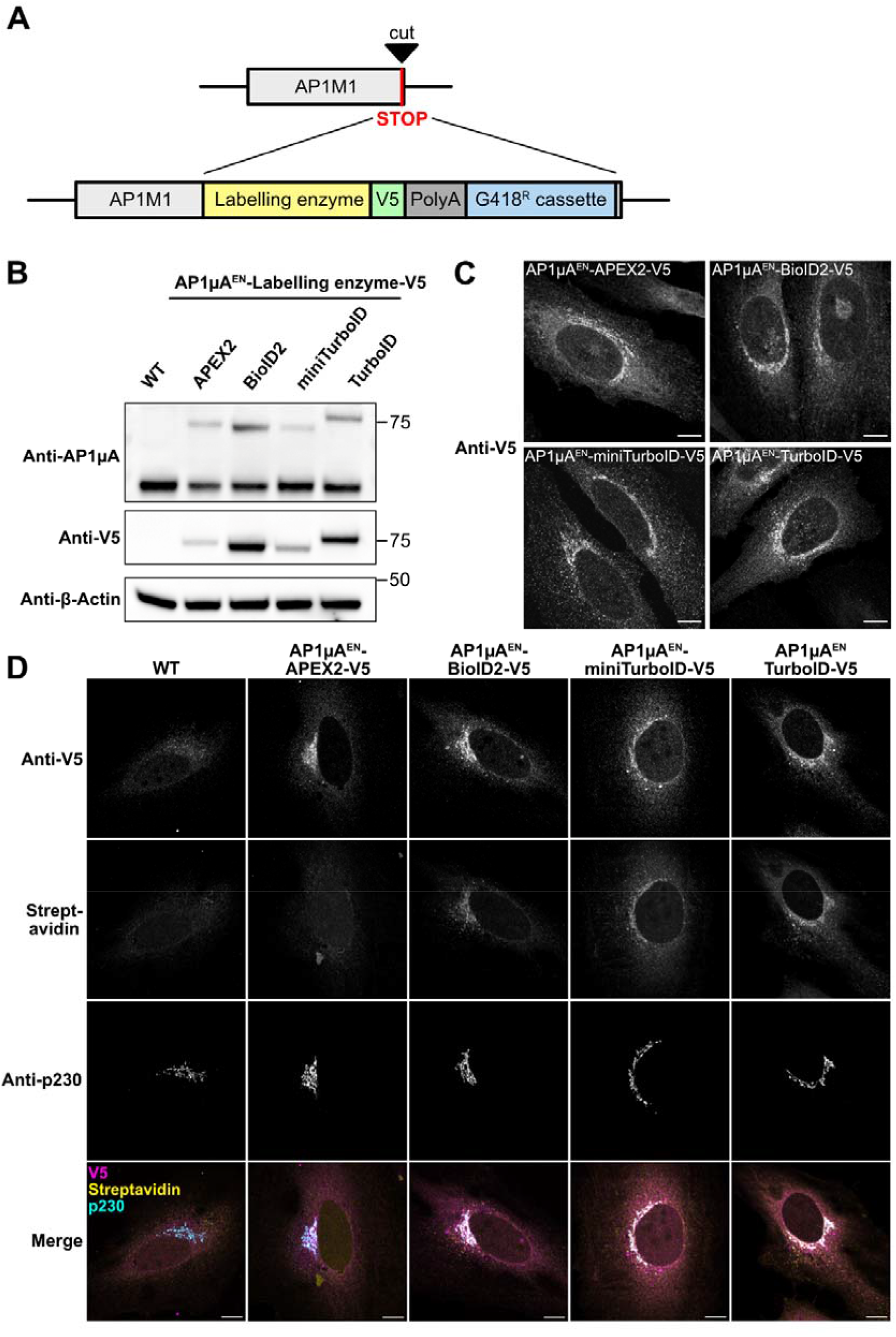
Rapid KI-strategy allows for endogenous tagging of AP1μA with different labelling enzyme for proximity biotinylation. **A:** Scheme of KI strategy. AP1μA was C-terminally tagged with the labelling enzyme, a V5 tag and a resistance cassette that allows for rapid selection of positive cells. **B:** Blots of whole cell lysates from generated cell lines to verify the KI of the labelling enzymes by anti-AP1μA and anti-V5 blotting. **C:** Cell lines expressing the labelling enzymes endogenously were fixed and stained for the V5 tag to detect the labelling enzyme expression and localization. **D:** Cells endogenously expressing the different labelling enzymes were fixed and stained with anti-V5 antibody to detect the labelling enzymes, anti-p230 antibody to mark the TGN area and streptavidin-AF488 to detect biotinylated proteins. Cells were treated with 50 μM biotin for 24 h and AP1μA^EN^-APEX2-V5-expressing cells were incubated for 30 min with 500 μM biotin-phenol and labelling was induced for 1 min with H_2_O_2_. ^EN^= endogenous. Scale bars are 10 μm.

Next, we wanted to compare the performance of the various labelling enzymes when expressed at endogenous levels. For biotin ligases, labelling times between less than 1h (miniTurboID and TurboID)^9,27^ and at least 16 h – 24 h (BioID2)^8,28^ are reported. To ensure the amount of biotinylation is sufficient for visualization and comparable between the different biotin ligases, we treated all cell lines with 50 μM biotin for 24 h to initiate the labelling. For the APEX2 peroxidase, we used the established effective labelling time of 1 min^6,29^ to avoid prolonged exposure to toxic H_2_O_2_. To visualize biotinylated proteins, we used fluorescently labelled streptavidin. The cargo adaptor complex AP-1 orchestrates transport between the TGN and endosomes^25,26,30^ and is reported to predominantly localize to the TGN^31^. Thus, we expected both the fusion protein and the pool of biotinylated proteins to localize in the TGN area. The different biotin ligases BioID2, miniTurboID and TurboID and the peroxidase APEX2 localize correctly when fused to endogenous AP1μA (Fig. 1D). Importantly, biotinylated proteins and AP1μA fusions are strongly concentrated to the TGN region, marked by the TGN resident protein p230^32^ (Fig. 1D). However, we could not find any specific biotinylation for the APEX2 peroxidase when expressed at physiological levels (Fig. 1D). To exclude general handling errors with the APEX2 sample, we transiently overexpressed Vimentin-APEX2 as well as AP1μA-APEX2 fusions and found for both constructs a specific biotinylation pattern (Supplementary Fig. 1A, B).

Quantitative analysis of the biotinylation rate of the BioID2, miniTurboID and TurboID KIs in immunofluorescence (Supplementary Fig. 1C, D) and western blot (Supplementary Fig. 1E, F) showed that shorter (2 h) labelling with MiniTurboID and TurboID leads to higher levels of biotinylated proteins than longer BioID2 labelling (24 h). This correlates well to what was originally reported for those labelling enzymes^8,9^. The low biotinylation rate of BioID2 makes it, in our opinion, unfavourable compared to the TurboID variants, as a high amount of biotinylation is crucial to enrich enough protein for the subsequent identification and quantification by mass spectrometry (MS). We decided to use the AP1μA^EN^-TurboID-V5 (^EN^= endogenous) cell line for our further experiments as the TurboID fusion yielded a higher biotinylation rate compared to the other fusions (Supplementary Fig. 1D).

### Physiological expression of TurboID fusions permits highly specific interactome mapping

Transient overexpression of a protein can lead to artefacts such as mislocalization or aggregation ^10-12^, increasing the chances of detecting non-specific, artificial interactors such as abundant cytosolic proteins. Physiological expression of the labelling enzyme should allow for highly specific, locally confined biotinylation of natural interactors. By applying two-color stimulated emission depletion (STED) super-resolution microscopy, we were able to visualize the biotinylation pattern after 2 h of biotin addition in AP1μA^EN^-TurboID-V5 knock-in cells (Fig. 2A) and compare it with AP1μA-TurboID-V5 overexpression (Fig. 2B). The high resolution achieved on the STED microscope enabled visualization of distinct nanodomains occupied by AP-1. Notably, the biotinylated proteins are primarily localized to the very same nanodomains as the endogenous AP1μA^EN^-TurboID-V5 fusion (indicated by the white arrows in Fig. 2A), indicating a high local specificity of the proximity labelling of the TurboID. Overall, localization of the overexpressed AP1μA-TurboID-V5 was more diffused (Fig. 2B) and we observed areas where biotinylated proteins and fusion protein did not overlap (white and yellow arrows in Fig 2B). Many cells overexpressing AP1μA-TurboID-V5 exhibit a very high background biotinylation in the cytoplasm and in the nucleus (Supplement Fig. 2A), while this was not noticed in the KI cells. Quantification of the background biotinylation revealed significantly higher cytosolic biotinylation in the transiently overexpressed cells compared to the KI cells, even after short biotin incubation (Supplement Fig. 2B).

**Fig. 2:**
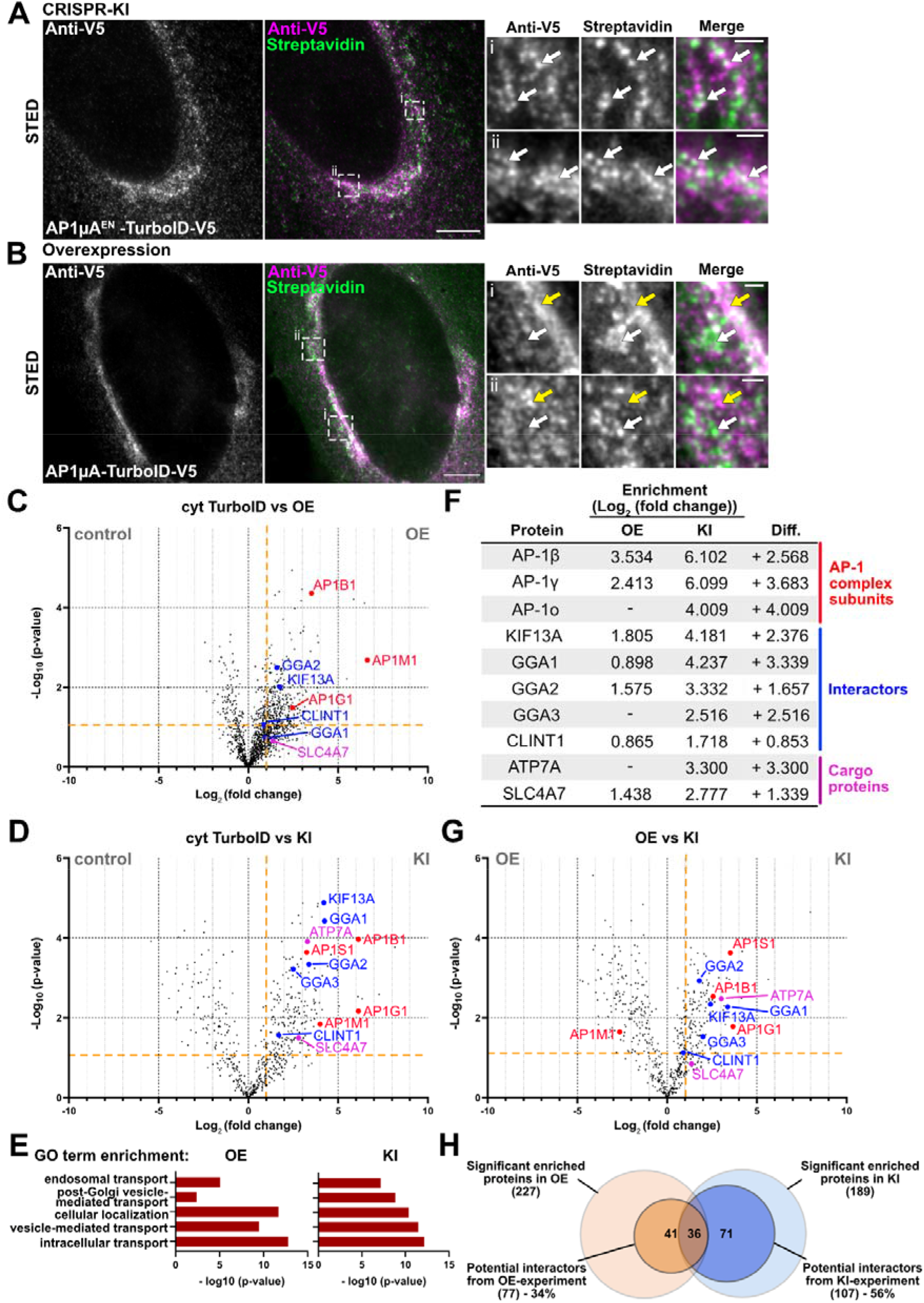
Endogenous tagging allows for more specific proximity labelling and interactome mapping than overexpression of the labelling enzyme. **A:** STED micrographs of a fixed AP1μA^EN^-TurboID-V5 cell stained with anti-V5 antibody and streptavidin-STARORANGE to detect biotinylated proteins. Cells were treated with 50 μM biotin for 2 h before fixation. Crops show distinct overlap of biotinylated proteins and AP1μA^EN^-TurboID-V5 (marked by white arrows). **B:** STED micrographs of a fixed cell transiently overexpressing AP1μA-TurboID-V5 that was treated as described in **A**. Crops show that biotinylated proteins and AP1μA-TurboID-V5 accumulate in distinct zones (white arrows indicate areas of biotinylation without AP1μA-TurboID-V5, yellow arrows indicate areas of accumulated biotin ligase without biotinylated proteins). Scale bars are 5 μm for STED images and 500 nm for crops in **A,B. C:** Volcano plot showing the changes in relative protein intensity between the overexpression (OE) experiment and control (cytosolic overexpressed TurboID). Significant hits are shown in the top right corner (p-value < 0.05 and log_2_ fold change > 1) separated by the orange lines. The volcano plot only includes proteins that were significantly enriched compared to WT cells. Subunits of the AP-1 complex (red), known interactors (blue) and known cargos (magenta) are marked. Entire protein list is shown in Supplementary Table 1. **D:** Volcano plot showing the changes in relative protein intensity between the KI experiment (AP1μA^EN^-TurboID-V5) and control. Same parameters as in **C. E:** GO term enrichment analysis showing enrichment of selected GO terms in overexpression (OE) and KI condition. **F**: Table of the analysed subunits, interactors and cargo proteins. Differences (Diff.) in log_2_ fold enrichment are indicated. **G:** Volcano plot showing the changes in relative protein intensity between KI experiment and OE experiment. Same parameters as in **C. H:** Venn diagram showing the number of potential interactors (defined by protein localization and function (see methods) and significant enrichment). Lists of potential interactors are shown in Supplementary Tables 2 and 3.

To probe whether endogenous tagging with TurboID allows biotin labelling of AP1μA-specific interactors with increased sensitivity compared to transient overexpression, we analyzed streptavidin-purified proteins by MS using label-free quantification. Biotin was added to the culture medium for 24 h to secure detectable protein labelling in the KI cells. We analyzed KI AP1μA^EN^-TurboID-V5 cells and cells transiently overexpressing an AP1μA-TurboID-V5 fusion protein. As controls we used cells transiently overexpressing a cytosolic TurboID-V5 as well as WT HeLa cells that were treated with biotin for 24 h. In total, we identified and quantified 4548 proteins (Supplementary Table 1). Proteins with at least a 2-fold increase in relative intensity compared to both controls (log_2_ fold change >1) and a p-value <0.05 were considered significantly enriched. We found 227 proteins significantly enriched in the overexpression sample and 189 proteins significantly enriched in the KI sample (Fig 2C,D). Overall, a larger number of proteins were enriched when AP1μA-TurboID was overexpressed compared to the KI condition, which is likely to be the result of mislocalization of AP1μA-TurboID-V5, possibly leading to the biotinylation of a larger set of proteins that naturally would not interact with AP-1. We found the two large subunits of the AP⍰1 complex (AP-1β1 and AP-1γ)^25^ significantly enriched in overexpression condition (Fig. 2C) and even more enriched in the KI cells (Fig. 2D). Notably, the other small subunit of the AP-1 complex (AP-1s)^25^ was only identified when AP1μA^EN^-TurboID-V5 was endogenously expressed. We next looked at known interactors of the AP-1 complex to test whether endogenous tagging improves enrichment of specific interactors. The AP-1 complex is recruited to the Golgi membranes where it directly binds to the clathrin adaptors EpsinR and GGAs (Golgi-localized, γ-ear-containing, Arf-binding proteins)^33,34^. Cell motility of membrane-bound AP-1 is conferred through the kinesin-like protein KIF13A, a microtubule-dependent motor protein that directly interacts with AP-1^35^. Most interactors were identified in the overexpression condition, but only GGA2 and KIF13A were found to be significantly enriched. Physiological expression of AP1μA^EN^-TurboID-V5, on the other hand, allowed significant enrichment of KIF13A, EpsinR (CLINT1) and all GGA proteins (GGA1-3). The difference between physiological expression levels and overexpression was even more striking when we looked at the AP-1-specific cargo proteins ATP7A and Sodium-bicarbonate co-transporter NBCn1 (SLC4A7)^36,37^. These AP-1 cargo proteins were all not or only slightly enriched in the overexpression experiment but significantly enriched in KI cells.

The overall impression that physiological expression of the TurboID fusion enhances the specificity of the hits we derive from our interactome data compared to overexpression can be confirmed when looking at gene ontology (GO) term analysis. While both data sets show enrichment of general GO terms such as *intracellular transport* or *vesicle-mediated transport*, AP-1-specific GO terms including *post-Golgi vesicle-mediated transport* or *endosomal transport*, are more enriched in the KI sample (Fig. 2E). Likewise, direct comparison between the two data sets shows significantly higher enrichment of all known interactors, subunits and cargo proteins in proximity labelling MS experiments performed in KI cells when compared to simple overexpression of the labelling enzyme (Fig 2F, G). The exception here is AP1μA itself, as the overexpression of the fusion proteins leads to higher enrichment in the overexpression (OE) condition (Fig. 2G).

To finally evaluate the overall quality of the MS data from the CRISPR-KI AP1μA^EN^-TurboID experiment, we identified possible interactors of AP1μA from the MS data and compared the two data sets (KI vs OE). We defined potential interactors as proteins that were significantly enriched and are either known to localize to the Golgi/TGN, are involved in cellular trafficking, are transmembrane proteins that might be trafficked by AP1μA or might be involved in the regulation of membrane homeostasis (e.g. regulatory kinases). For the KI cell line, we found a total of 107 potential interactors (Supplementary Table 2), which corresponds to 56% of all significant enriched hits in the MS. Evaluation of the MS data from the overexpressed AP1μA-TurboID resulted in a list of only 77 potential interactors (34% of total number of significant hits) (Supplementary Table 3). The results of our analysis are illustrated in Fig. 2H. Taken together, our findings highlight that tagging the protein with the labelling enzyme endogenously increases the sensitivity of the proximity biotinylation and the MS measurement so that very transient but specific interactors, such as specific cargo proteins, can be identified.

### Interactome mapping of different AP-complexes with endogenous TurboID fusions

The comprehensive interactome data resulting from endogenous tagging of AP1μA with TurboID encouraged us to apply the rapid knock-in TurboID approach to different proteins to probe its versatility and, in particular, to reveal the native interactome of AP complexes. Five different AP complexes (AP-1, AP-2, AP-3 and AP-4 and the more recently discovered AP-5) are responsible for sorting cargo throughout the endo-lysosomal system of human cells^38^. Our overall understanding of the intracellular role of the different AP complexes would greatly benefit from a better knowledge of their interactome. Aside from a few cargo proteins that are often used as model cargos, little is known about which proteins are sorted by which adaptor protein in mammalian cells^39^. Recent proteomic studies have shed light on cargo sorting pathways in yeast cells^40^, however such a comparative and comprehensive study is still lacking for mammalian cells. We also do not fully understand the mechanisms of recruitment of AP complexes to different membranes. Although AP-1, AP-3 and AP-4 are all known to be recruited by ARF1 to TGN/endosomal membranes^41^, they localize to distinct endosomal buds^42,43^, suggesting the presence of different unknown interactors. Due to their role as key regulators of intracellular trafficking, dysfunction of AP complexes is linked to a variety of diseases^38,41,44^ and a better understanding of the interaction partners could help reveal the underlying mechanistic basis of pathology. While AP-1 is known to locate to the TGN and endosomal membranes^26,43,45^, where it coordinates clathrin-dependent trafficking between the two organelles^25,46,47^, AP-2 is found at the plasma membrane, where it recruits clathrin for clathrin-mediated endocytosis^25^. AP-3 is thought to localize to the same endosomes as AP-1 but has different cargo clients, which points towards a role in trafficking to the lysosome^43,48^. AP-4 binds to TGN membranes^38^, where it mediates transport of autophagosomal factor ATG9A^49,50^. The low abundance of AP-4 (about 40 fold lower than AP-1 or AP-2 in HeLa cells^51^) makes it an interesting target to test the endogenous TurboID tagging on a very low abundant protein. AP-5 is thought to be involved in the transport from the late endosome to the Golgi or the lysosome and to also contribute to lysosome maintenance^52,53^. It is different from other AP complexes as it associates with two additional proteins^54^ and is not recruited to intracellular membranes by ARF1 as are AP-1, AP-3 and AP-4. However, as it is as comparably low abundant as AP-4^53^ and was not precisely localized so far, we decided not to include it in this study as it would be difficult to confirm correct integration of the endogenously tagged subunit.

We C-terminally tagged the μ-subunit of the different AP complexes with TurboID and tested the expression and localization of the fusion protein with immunofluorescence imaging and western blot (Fig. 3A,B). After 24 h of biotin treatment, we were able to detect extensive biotinylation of proteins via western blot and in immunofluorescence imaging experiments for all AP complexes, even for the low abundant AP4μ^EN^-TurboID-V5 fusion. To map and compare the interactome of the four AP complexes we analysed streptavidin-purified biotinylated proteins by MS using label-free quantification for all the KIs. As a control we used HeLa WT cells that were treated with biotin. In total, we identified and quantified 2574 proteins (Supplementary Table 4). In order to identify specific interactors and cargo proteins for each AP complex, we compared the relative intensity of a given protein between data sets for various AP complexes. Proteins with at least a 2-fold increase in relative intensity (log_2_ fold change >1 or <-1) and a p-value <0.05 were considered significantly enriched. By doing so, we were able to identify AP-2 complex specific interactors such as Epsin-1 (EPS1) and Epsin-2 (EPS2), Synaptojanin-1 (SYNJ1) and the protein numb like (NUMB) that are all known for their role in clathrin mediated endocytosis (Fig. 3C). Importantly, the data set for the low abundant AP4μ^EN^-TurboID-V5, highlighted several known AP-4 specific interactors such as Tepsin (ENTDH2), Hook1 (HOOK1) and a FHF complex subunit (FAM160A1)^55^ (Fig. 3D). Comparison of the interactome of various AP complexes allowed not only the identification of interactors that are specific for one single AP complex (e.g. GGA proteins and the PI4-kinase PI4K2B for AP-1^56,57^), but also revealed potential common interactors such as the SNARE protein Vamp7 that is enriched for AP-1 as well as AP-3 (Fig. 3D). Aside from potential uncharacterized effectors, endogenous TurboID-tagging enabled the identification of potential cargo proteins specific for each AP complex. We identified known cargo proteins (various integrins (ITGA5, ITGAV or ITGA1) for AP-2 or ATG9A for AP-4) and also novel potential cargo proteins such as the ring finger protein 121 (RNF121), a Golgi localized protein with anti-apoptotic effects in cancer cells^58^, the plasma-membrane-localized cation channel TRPM7, the TGN-localized calcium transporter ATP2C2 for AP-1 and the lysosomal Cl^-^/H^+^ Antiporter ClC-7 (CLCN7) for AP-3 (Fig. 3C,D). Those potential cargo proteins all have at least one tyrosine-based YXXφ motif in their cytoplasmic domains that is one of the motifs necessary for sorting via AP-1 or AP-3. We compared the significantly enriched proteins from the data sets of each AP complex and assembled a list of potential interactors and potential cargo proteins for each AP complex (Supplementary Table 5). Potential interactors and cargo proteins were defined proteins that are known to be involved in membrane trafficking, might be involved in regulation of membrane homeostasis (e.g. regulatory phosphatases and kinases), have a transmembrane domain (potential cargo proteins) or are known to play a role in clathrin mediated endocytosis (for AP-2). In total, we identified more than 300 potential interactors and more than 200 potential cargo proteins.

**Fig. 3:**
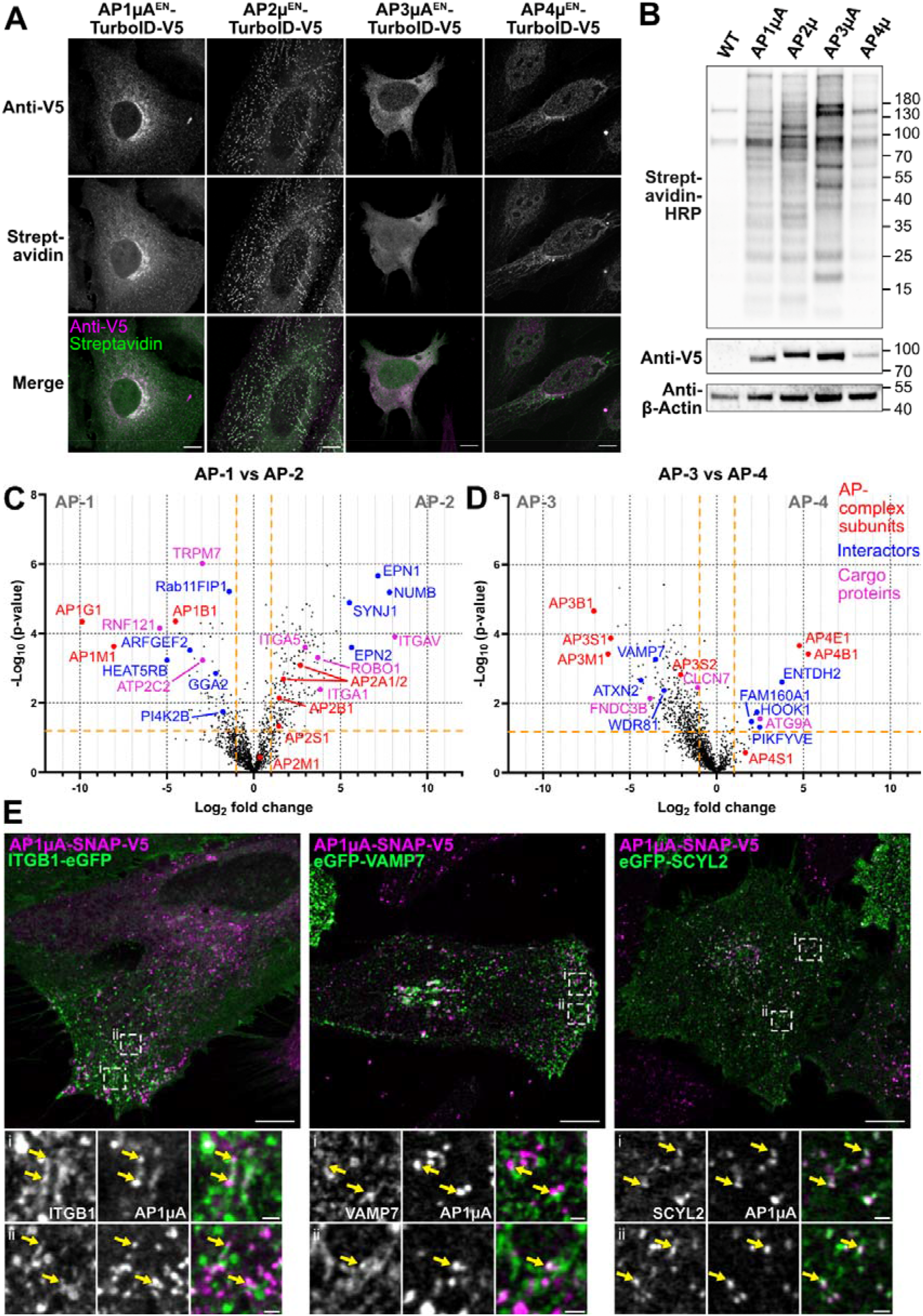
Interactome comparison of AP complexes with endogenous TurboID-tagging. **A:** HeLa KI cells expressing the endogenous AP μ subunits 1-4 fused to TurboID-V5 were fixed and stained with anti-V5 antibody to detect the labelling enzyme and streptavidin-AF488 to detect biotinylated proteins. Cells were incubated with biotin (50 μM) for 24 h before fixation. **B:** Visualization of TurboID activity in all four KI cell lines on a western blot. Cells were treated with 50 μM biotin for 24 h. Whole cell lysates were blotted with streptavidin-HRP to detect biotinylated proteins, and anti-V5 antibody to detect ligase expression. **C:** Volcano plot showing the changes in relative protein intensity between AP1μA^EN^-TurboID-V5 and AP2μ^EN^-TurboID-V5. Proteins that show significant changes in their relative intensity are shown in the top left (AP-1) and top right (AP-2) corner (p-value <0.05 and log_2_ fold change >1 or <-1) separated by the orange lines. Subunits of the AP complexes (red), potential interactors (blue) and potential cargos (magenta) are marked. **D:** Volcano plot showing the changes in relative protein intensity between AP3μA^EN^-TurboID-V5 and AP4μ^EN^-TurboID-V5. Same parameters as in **C. E:** HeLa AP1μA^EN^-SNAP-V5 KI cells labelled with JFX650-BG that were transiently transfected with plasmids encoding for ITGB1-eGFP, eGFP-VAMP7 and eGFP-SCYL2 (left to right). Crops show where AP1μA domains are observed in close proximity to structures defined by the various proteins tested (marked by yellow arrows). Scale bars are 10 μm and 1 μm for crops in **A,E**.

We then further tested some of the unexpected hits we found in the AP1μA data. We selected Integrin beta 1 (ITGB1), the SNARE protein VAMP7 and SCY1-like 2 (SCYL2). ITGB1 could be a potential cargo protein of AP-1^59^. VAMP7 is a component of a SNARE complex composed of syntaxin-8, syntaxin-7, VAMP7 and VTI1B that is involved in endosomal recycling of endocytosed material^60^, and interestingly, we found all four members as potential AP-1 interactors. So far, VAMP7 was only reported to interact with AP-3, and no direct interaction with the other adaptor complex, AP-1, was observed^61,62^. SCYL2 was originally identified as a protein kinase for AP-2^63^ but has also been connected to AP-1- and AP-3-mediated trafficking^64^ as well as clathrin-dependent TGN export in plants^65^, but its overall role remains poorly understood. We transiently expressed GFP-fusions of the three proteins in an AP1μA^EN^-SNAP-V5 KI cell line, where AP1μA has been tagged with a SNAP tag (Supplementary Fig. 3), enabling visualization of endogenous AP1μA in living cells. Live-cell confocal microscopy showed that vesicular AP-1 structures can be found on tubular compartments positive for either ITGB1 or VAMP7 (Fig. 3E). Similarly, we find punctuated SCYL2 structures perfectly colocalizing with vesicular AP1μA (Fig. 3E). All three proteins are thus likely to interact with the adaptor complex AP-1, as suggested by their close proximity in living cells. Interestingly, ITGB1 is found together with AP-1 in the peripheral areas of the cell, pointing towards a possible role for AP-1 in the endocytic recycling of the integrin, a role that has been described for AP-1 in the recycling of the transferrin receptor^66^. The close proximity of SCYL2 and VAMP7 to vesicular AP-1-positive structures suggests some regulatory functions of those proteins in AP-1 mediated trafficking.

### N-terminal endogenous tagging to map the interactome of CLCa

To test whether we can apply our rapid KI strategy also for N-terminally tagged proteins, we fused the different labelling enzymes to the N-terminus of endogenous clathrin light chain A (CLCa). To create N-terminal fusions, the resistance cassette is inserted between LoxP sites upstream of the start codon of the V5-tag and the TurboID^18^. In a second step, the resistance cassette is then excised via transfection of Cre recombinase. This step becomes necessary as the presence of the resistance cassette in the genome could possibly isolate the gene from its promoter region and cause it not be transcribed. The strategy used for N-terminal tagging is illustrated in Fig. 4A. Successful expression of the fusion protein was confirmed on western blot and with immunofluorescence microscopy (Fig. 4B, C). Moreover, immunofluorescence microscopy of the CLCa-fusion proteins together with clathrin heavy chain (CHC) shows that both proteins colocalize, suggesting correct localization and function of the fusion proteins (Fig. 4D). The overall KI rates were comparable to C-terminal tagging (55% (APEX2), 25% (BioID2), 51% (MiniTurboID), 71% (TurboID)). Alike for C-terminally tagged AP1μA, we observed locally confined biotinylation for the biotin ligases but no specific biotinylation for the APEX2 fusion protein (Fig. 4D).

**Fig. 4:**
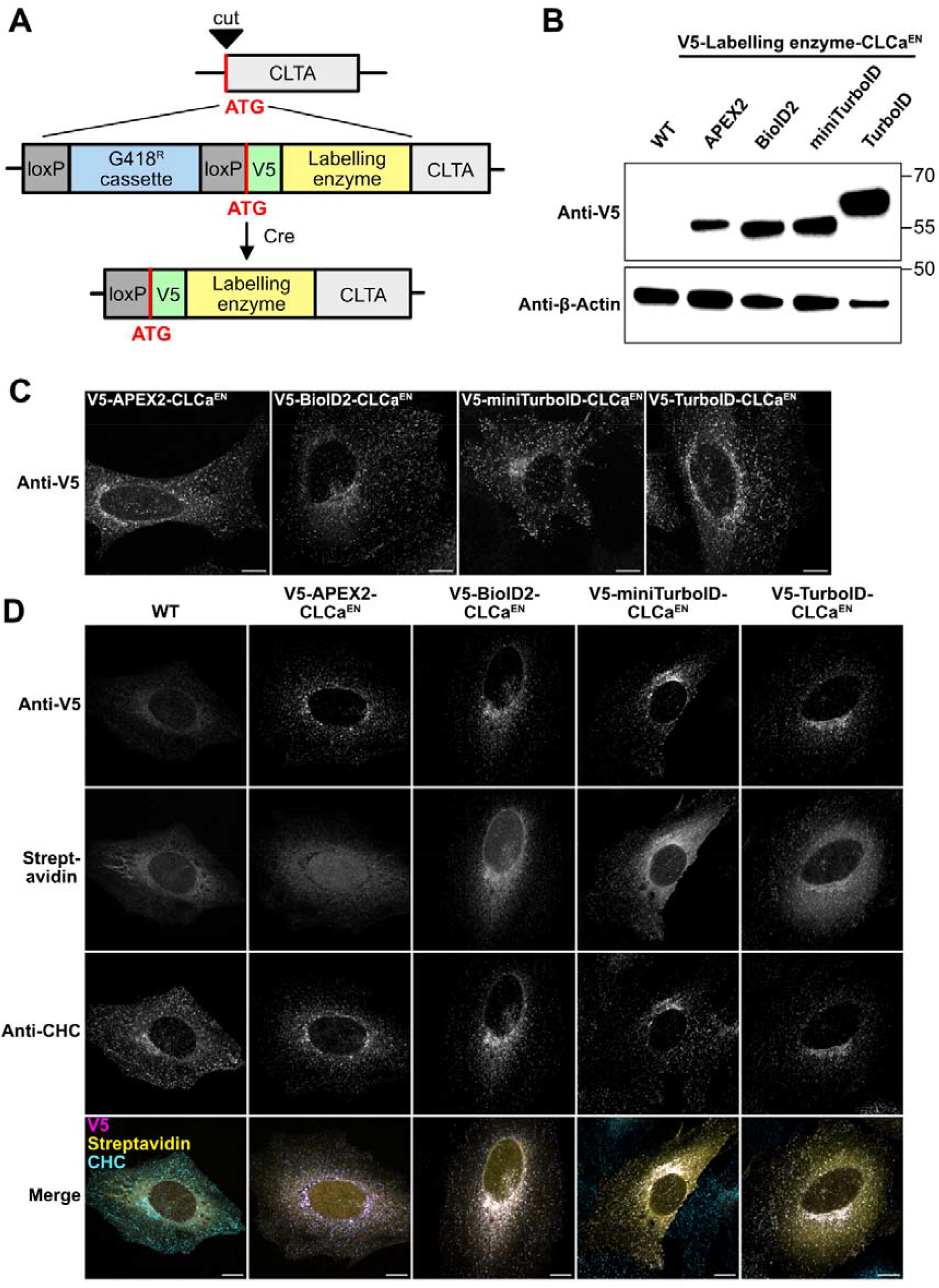
Endogenous N-terminal tagging of CLCa with labelling enzymes. **A:** Scheme of KI strategy. CLCa was N-terminally tagged with the labelling enzyme and a V5 tag. The integrated resistance cassette can be excised with the loxP-Cre-system after transfection with the Cre recombinase. **B:** Blots of whole cell lysates from generated cell lines to verify the KI of the labelling enzymes by anti-V5 blotting. **C:** Cell lines expressing the labelling enzymes endogenously were fixed and stained for the V5 tag to detect the labelling enzyme expression. **D:** Cells endogenously expressing the different labelling enzymes were fixed and stained with anti-V5 antibody to detect the labelling enzymes, anti-CHC antibody to detect the endogenous clathrin and streptavidin-AF488 to detect biotinylated proteins. Cells were treated with 50 μM biotin for 24 h and V5-APEX2-CLCa^EN^ expressing cells were incubated for 30 min with 500 μM biotin-phenol and labelling was induced for 1 min with H2O2. Scale bars are 10 μm.

We then used the V5-TurboID-CLCa^EN^ KI cell line to map the CLCa interactome by analyzing streptavidin-purified biotinylated proteins with MS using label-free quantification (Supplementary Table 6). CLCa fulfils multiple roles in intracellular trafficking such as coating membranes during endocytic events or trafficking intermediates shuttling between the Golgi/TGN and endosomes^67^. Clathrin does not directly bind membranes but uses adaptor proteins such as adaptor protein complexes AP-1 for Golgi-endosome trafficking and AP-2 for clathrin-mediated endocytosis^25^. As we already have performed interactome mapping experiments for those two adaptors, a direct comparison between the data sets generated from CLCa and AP-1 or AP-2 TurboID KIs allows the sequestering of CLCa interactors that are involved in endocytosis from those involved in intracellular post-Golgi transport. A comparison of CLCa against AP-1 shows strong enrichment of proteins such as Epsin-1 (EPS1), Synaptojanin-1 (SYNJ1) and the protein numb like (NUMB) that are known for their role in clathrin-mediated endocytosis (Fig. 5A). Furthermore, GO terms such as *endocytosis, clathrin-mediated endocytosis* and *import into cell* are strongly enriched (Fig. 5B). On the contrary, comparison of the interactome of CLCa with AP-2 shows proteins enriched that are involved in AP-1 dependent post-Golgi transport such as the GGA proteins and the HEAT repeat containing 5B (HEATR5B)^66^ (Fig. 5C). Here, among the most enriched GO terms we find *intracellular transport* and *Golgi-vesicle transport* (Fig. 5D). In conclusion, endogenous N-terminal tagging of CLCa with TurboID granted a highly specific interactome data set that, when combined with data sets from AP-1 or AP-2, allowed mapping of pathway-specific interactors.

**Fig. 5:**
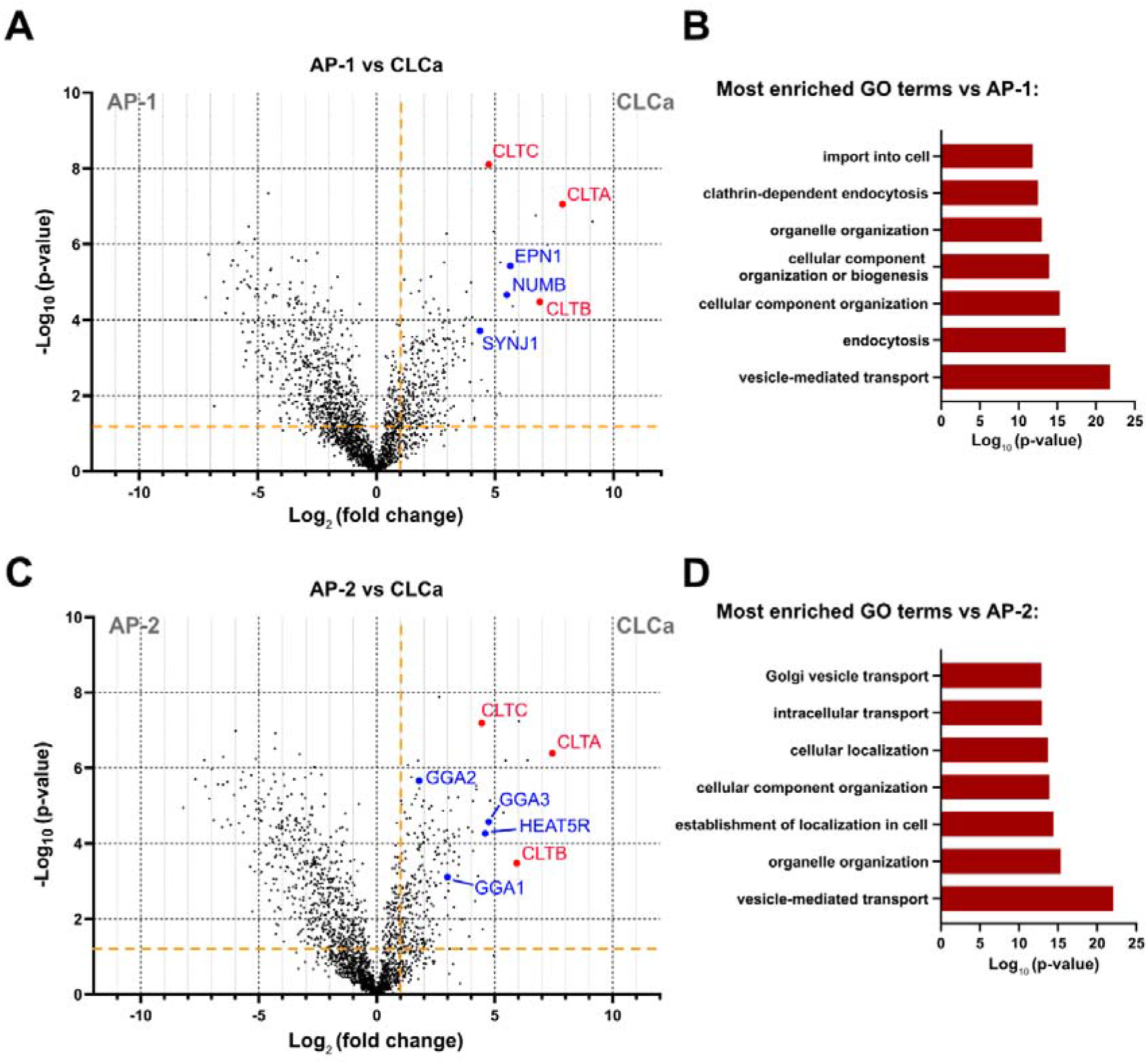
Comparison of interactome data set allows mapping of pathway specific clathrin interactors. **A:** Volcano plot showing the changes in relative protein intensity between AP1μA^EN^-TurboID-V5 and V5-TurboID-CLCa^EN^. Significant hits (p-value <0.05 and log_2_ fold change >1) are separated by the orange lines in the top right. Clathrin chains (red) and exemplary proteins involved in clathrin-mediated endocytosis (blue) are marked. **B:** GO term enrichment analysis showing the most enriched GO terms comparing enrichment for CLCa against AP1μA. **C:** Volcano plot showing the changes in relative protein intensity between AP2μ^EN^-TurboID-V5 and V5-TurboID-CLCa^EN^. Same parameters as in **A**. Clathrin chains (red) and exemplary proteins involved in post-Golgi transport (blue) are marked. **D:** GO term enrichment analysis showing the most enriched GO terms comparing enrichment for CLCa against AP2μ.

## Discussion

Employing an antibiotic-based CRISPR KI strategy for C- and N-terminal tagging, we endogenously fused the commonly used labelling enzymes APEX2, BioID2, miniTurboID and TurboID to AP1μA and CLCa (Fig. 1, 4). We identified the biotin ligase TurboID as best suited for the KI approach in combination with proximity labelling experiments, as it combines high KI rates with favourable labelling kinetics (Supplementary Fig. 1). The lower KI rates of miniTurboID compared to TurboID might be result of lower stability of miniTurboID fusion proteins, which has already been reported in other studies^9,68^. Faster labelling kinetics are not only beneficial to reliably generate sufficient amounts of biotinylation to enrich enough protein for the subsequent identification and quantification by MS but are also advantageous when designing experiments that require short labelling times. Generally, the incubation time with the biotin might be adapted according to the abundance of the POI, when working with highly abundant proteins, incubation times shorter than 24 h with biotin could be used. Previous work has already demonstrated the TurboID variants outperform BioID or BioID2^9,27,28,68^, however BioID2 is still a commonly used enzyme. Recently, new variants of the BioID2 were introduced, microID and ultraID^69^. With a molecular weight below 20 kDa, they are significantly smaller than the here presented labelling enzymes. Especially the ultraID is reported to have labelling kinetics similar to TurboID with less background activity. In addition, an ancestral BioID variant called AirID was developed with faster labelling kinetics than BioID and more specific labelling compared to TurboID^70^. Another possibility to reduce labelling background of TurboID is the use of a light-controlled variant of the TurboID (LOV-Turbo)^71^. It would be interesting to see how these variants perform when used for tagging at the endogenous locus. Practically, to prevent background biotinylation of TurboID, cells can be grown in biotin-free media 72 h prior to the experiment. This becomes crucial in interactome mapping experiments that, for example, require induced changes of the cell state, as labelling should only occur after the change was induced. To our surprise, we could not achieve any biotinylation with the APEX2 peroxidase expressed as a low abundance endogenous fusion (Fig. 1, 4). The short labelling time in combination with the low physiological expression does not yield detectable proximity labelling in our hands. Generally, CRISPR KI approaches to tag endogenously with APEX2 were reported to be functional^15,72^; however, for us, multiple attempts to induce labelling failed. We can only speculate that highly concentrated APEX2 is needed to get sufficient biotinylation when it is endogenously expressed. Specific confined cellular environments, such as nucleus or stress granules, would favour such a high concentration of an APEX2 fusion protein. On the other hand, soluble cytosolic proteins may not be optimal for an APEX2 approach. Yet, labelling with TurboID and biotin worked reliably and was easy to induce for peripheral cytoplasmic machinery like adaptors and clathrin.

Using the AP1μA^EN^-TurboID-V5 cell line created with the KI strategy, we were able to compare the enrichment of known interactors and cargo proteins of the AP-1 complex in quantitative MS measurements against data sets generated with cells overexpressing AP1μA-TurboID-V5. Strikingly, we found known interactors to be significantly more enriched when AP1μA^EN^-TurboID-V5 was expressed at endogenous levels (Fig. 2). Especially relevant for our research is the strong enrichment of AP-1 cargo proteins that can be only observed when TurboID is endogenously fused to AP1μA, highlighting the importance of matching the endogenous expression levels. The increased sensitivity for real interactors is likely a result of less unspecific labelling due to mislocalization- or aggregation artefacts induced by overexpression of the labelling enzyme fused to the POI. A larger background of peptides from unspecifically labelled proteins increases the sample complexity and therefore lowers the overall sensitivity of the MS measurements, as all peptides compete for ionization and detection. Endogenous protein tagging with a biotin ligase that allows for highly confined biotinylation (Fig. 2A), therefore increases the chances for detection of specific interactors, especially if they are of low abundance. Importantly, endogenous expression of the AP1μA^EN^-TurboID fusion resulted in a more comprehensive list of potential interaction partners compared to overexpression of AP1μA-TurboID (Supplementary Tables 2, 3 + Fig. 2E,H). Using the described KI-pipeline, we tagged the μ-subunit of different AP complexes with TurboID and comparably analyzed their interactome in quantitative MS measurements (Fig. 3). The provided lists of potential interactors and cargo proteins for each individual AP complex (Supplementary Table 5) not only demonstrate the versatility of the approach but also present a database that can contribute to the better understanding of AP-driven intracellular sorting. Our data sets can be used as a starting point for studies aimed at unravelling the mechanisms of spatially confined recruitment of different TGN- and endosome-associated AP complexes. Furthermore, linking the AP complexes to the different potential cargo proteins will shine light on the intracellular sorting and trafficking routes of those proteins. By picking three proteins from our list of potential AP-1 interactors (ITGB1, VAMP7 and SCYL2) and studying their localization in respect to AP-1 in living cells (Fig. 3E), we could show that they all localize in the vicinity of AP-1. Lastly, we mapped the interactome of the N-terminally TurboID-tagged CLCa and compared the data to the data sets of AP1μA and AP2μ (Fig 5, Supplementary Table 6). By doing so, we were able to identify pathway-specific interactors of CLCa.

In summary, endogenous tagging with biotin ligases enables highly specific proximity labelling and increases the sensitivity for real interactors that might be transient or of low abundance. Our pipeline presents an alternative to classical CRISPR-based approaches that would require single-cell selection of positive TurboID clones and can be applied to any cell line as long as it can be reliably transfected and selected. The pipeline for the generation of CRISPR KIs, based on the integration of a resistance cassette, allows rapid generation of endogenously tagged cells and can be applied to low abundant target proteins like AP-4. Aside from obvious time and work reduction, it also avoids artefacts that arise from single clone behaviour. In particular, for naturally low abundant proteins, inducible systems may still trigger non-physiological protein expression.

Rapid endogenous tagging with TurboID enabled us to map and compare the interactome of 4 different AP complexes and revealed known but also novel AP specific interactors. The ease and speed of the KI generation makes it an attractive alternative to transient overexpression of the labelling enzyme or classical KI approaches using single cell selection, as the MS experiments can be done in about 4-5 weeks after the CRISPR transfection. Additionally, the tools provided here can be used in a variety of different cell lines for the identification of cell type-specific and physiological cargoes.

## Acknowledgments

We acknowledge the whole lab for the fruitful discussion during the planning and writing phase of this manuscript. This project was supported by Deutsche Forschungsgemeinschaft (DFG, German Research Foundation) grants SFB958 (Project A25), SFB/TRR186 (Project A05 (CF) and Project A20 (FB)) and a major research instrumentation grant for the acquisition of the STED microscope. For mass spectrometry, we acknowledge the assistance of the Core Facility BioSupraMol supported by the Deutsche Forschungsgemeinschaft (DFG).

## Methods

### Antibodies and Streptavidin conjugates

All dyes and antibodies used in this study are provided in Supplementary Table 7 in the supplementary information.

### Mammalian cell culture

All experiments were carried out in HeLa cells ATCC CCL-2 (ECACC General Collection) grown in a humidified incubator at 37°C with 5% CO_2_ in DMEM (Gibco-ThermoFisher) supplemented with 10% fetal bovine serum (Corning) and penicillin/streptomycin (Lonza Bioscience) to prevent contamination. For transient transfection, HeLa cells at 70-80% confluency were transfected with FuGENE HD Transfection Reagent (Promega) according to the supplier’s protocol.

### Generation of CRISPR-Cas9 knock-in cell lines

See supplementary information.

### Plasmid design of overexpression plasmids

See supplementary information.

### Immunofluorescence

For all immunofluorescence samples, 40 000 cells were seeded on fibronectin-coated coverslips. Biotinylation was induced by replacing the growth media with medium containing 50 μM biotin and samples were incubated for 24 h or 2 h as indicated in the figures. Biotinylation for AP1μA^EN^-APEX2-V5 cells was induced as described by Hung et al.^29^. All cells were washed twice with PBS and then fixed in 4% PFA for 10 min at RT. Subsequently, they were rinsed three times with PBS and incubated for 3 min in permeabilization buffer (0.3% NP40 (Roth), 0.05% Triton-X 100 (Sigma Aldrich) and 0.1% BSA (IgG free) (Roth) in PBS). Cells were blocked for 1 h in blocking buffer (0.05% NP40, 0.05% Triton-X 100 and 5% goat serum (Jackson ImmunoResearch) in PBS) at RT and then incubated with primary antibodies in blocking buffer overnight rocking at 4°C. On the next day, the samples were washed three times 5 min in washing buffer (0.05% NP40, 0.05% Triton-X 100 and 0.2% BSA (IgG free) in PBS) before incubation with the respective secondary antibodies in blocking buffer for 1 h rocking at RT. For visualization of biotinylated proteins either Streptavidin-Alexa488 (for confocal microscopy) or Streptavidin-STARORANGE (for STED microscopy) was added to the secondary antibody mix. The cells were then washed three times 5 minutes with wash buffer and then dipped in dH_2_O before mounting with ProLong Gold Antifade Reagent (Thermo Fisher Scientific). Mounted samples were let to harden overnight at RT and then stored at 4°C until imaging.

### Live-cell imaging

For live-cell imaging experiments, 100.000 cells were seeded on glass-bottom dishes (3.5 cm diameter, No. 1.5 glass; Cellvis), coated with fibronectin (Sigma) beforehand. One day after seeding, KI cells expressing the AP1μA-SNAP fusion were labelled using an O^6^-benzylguanine (BG) substrate (JFX650-BG) (1μM) in culture medium for 1 h^73^. After the labelling, cells were washed for at least 1 h in culture medium. Live-cell imaging was carried out in FluoroBrite DMEM (Gibco) supplemented with 10% FBS, 20 mM HEPES (Gibco) and GlutaMAX (Gibco). For live-cell imaging experiments a microscope incubator (Okolab) was used to keep the stage and sample at 37°C.

### Imaging and image processing

Confocal and STED imaging was carried out on a commercial expert line Abberior STED microscope equipped with 485 nm, 561 nm and 645 nm excitation lasers. For two-color confocal live-cell imaging both signals were detected simultaneously, detection windows were set to 498 to 551 nm and 650 to 756 nm. For two-color STED experiments both dyes were depleted with a 775 nm depletion laser. The detection windows for the dyes were set to 498 to 551 nm, 571 to 630 nm and 650 to 756 nm. Excitation power was kept constant between samples in the same experiment to be able to quantify differences in expression levels and biotinylation. The pixel size was set to 60 nm for confocal and 20 nm for STED.

All images shown were smoothed using a Gaussian filter with 1-pixel SD using ImageJ ^74^. For better representation of the AP4μ^EN^-TurboID-V5 and live-cell images, the background was subtracted using a rolling ball radius of 50.0 pixels.

### Image analysis and statistical analysis

All image analysis was carried out with ImageJ. To determine the ratio of biotinylated proteins and V5-tagged AP1μA-labelling enzyme fusions in Supplementary Fig. 1D, a small region in the Golgi area was selected in the raw image and the average grey values were measured for both channels. The ratio between the mean intensity fluorescence of the biotinylated proteins in the Golgi area versus the mean fluorescence intensities from the V5 channel was then calculated. For each condition at least 40 cells were analysed from three independent experiments.

To analyse background biotinylation in Supplement Fig. 2B, a small region in a cytoplasm was selected and the mean intensity fluorescence was measured. At least 20 cells from each condition were analysed.

Statistical analysis (t-tests) was carried out with Prism. P-values are indicated in figure legends.

### Quantification of KI rate

To estimate the number of gene-edited cells, V5-immunostained cells for each KI cell line were screened for the presence of V5 signal using confocal light microscopy. 100 cells were screened for each cell line.

### Knock-in verification via western blot

For each knock-in cell line, 180 000 cells were seeded on a 6-well plate. 24 h later cells were washed twice with PBS and harvested in 350 μl of Laemmli sample buffer. The samples were boiled for 10 min at 98°C before loading 30 μl of each sample on two separate 4-12% SDS-polyacrylamide gels. After electrophoresis, proteins were transferred to a nitrocellulose membrane (Amersham) via wet blotting. Both membranes were washed once with phosphate buffered saline with 0.05% Tween (PBST) and then blocked for 1 h with 5% (wt/vol) milk powder and 1% BSA in PBST rocking at RT. After that, membranes were washed once 5 min with PBST and two times 5 min with PBS before incubation with the respective antibodies overnight rocking at 4°C. On the next day, the membranes were washed three times 5 min with PBST and incubated with a secondary antibody coupled to HRP in 5% (wt/vol) milk powder and 1% BSA in PBST for 30 min rocking at RT. The membrane was washed twice with PBST and twice with PBS for 5 min each. To develop the membrane the ECL western blot substrate was added for 2 min and then the membrane was imaged.

### Western blot analysis of biotinylated proteins

For detection of biotinylated proteins on a western blot, 800 000 HeLa WT and all KI cell lines were seeded on a 10 cm cell culture dish. Starting on the next day, the media was replaced with biotin-containing medium (50 μM biotin) and samples were incubated for 24 h or 2 h at 37°C. The biotin addition was timed in a way that all samples could be harvested at the same time. Biotinylation for AP1μA^EN^-APEX2-V5 cells was induced as described Hung et al.^29^. For harvesting, the cells were washed twice with ice-cold PBS and then extracted in 400 μL of ice-cold lysis buffer (50 mM Tris pH 7.4, 150 mM NaCl, 2 mM EDTA, 0.5% NP-40, 0.5 mM DTT and protease inhibitors). Cells were then centrifuged for 10 min at 4°C at 14,000g to clarify the cell lysates. 20 μl of the whole-cell lysates were mixed with Laemmli buffer and boiled at 95°C for 10 min before loading on a 4-12% SDS-polyacrylamide gel. Two separate gels were used, one for detection of biotinylated proteins and the other for detection of the fusion protein. After electrophoresis, proteins were transferred to a nitrocellulose membrane (Amersham) via wet blotting. To visualize biotinylated proteins on the membrane, after blocking (5% (wt/vol) milk powder in PBST), the blot was incubated with 0.3 μg/mL streptavidin-HRP in 3% BSA in PBST for 30 min rocking at RT. The western blot to detect the fusion protein with the V5 tag was carried about as described above.

For quantification of the biotinylated proteins in relation to the available labelling enzyme (Supplementary Fig. 1F) the intensity of the line on the biotinylation blot was measured in ImageJ and set in relation to the intensity of the corresponding V5 line. To measure the intensity of a line of biotinylated proteins in ImageJ, a box was drawn in the middle of the measured signal and the average intensity value of this area was used for analysis. The respective position and size of the box was kept constant for each line of a blot. The V5 tag intensity was measured accordingly except that the box was always placed around the area of the strongest signal. For each experiment the values were normalized to the AP1μA^EN^-TurboID-V5 sample that was incubated for 24 h with biotin. The experiment was repeated three times.

### Preparation of MS samples, LC-MS and data analysis

See supplementary information.

### Identification of potential interactors of AP1μA comparing KI and overexpression

For identification of potential interactors only proteins that were significantly enriched (p-value <0.05 and log_2_ fold change > 1) against both WT cells and cytosolic TurboID control were considered. These proteins were then screened with Uniprot to identify proteins that are either known to localize to either the Golgi or TGN, are thought to be involved in cellular trafficking, or are transmembrane proteins that might be trafficked by AP1μA or could control trafficking events (e.g. kinases, phosphatases). Together with all uncharacterized proteins these proteins were then considered potential interactors and summarized in Supplementary Tables 2,3.

### Identification of potential interactors and cargo proteins comparing different AP complexes

For identification of potential interactors and cargo proteins only proteins that were significantly enriched (p-value <0.05 and log_2_ fold change > 1) compared to at least one other AP-complex and also higher enriched than in the WT control (log_2_ fold change > 0) were taken into consideration. These proteins were then screened with Uniprot to identify proteins that are either known to localize to either the Golgi and endosomal system (or to CCP in the case of AP-2), are thought to be involved in cellular trafficking, or are transmembrane proteins that are potential cargo proteins or could control trafficking events (e.g. kinases, phosphatases). All of those proteins that have at least one transmembrane domain were considered potential cargo proteins, all others were termed potential interactors. The results were summarized in Supplementary Table 5 (switch between sheets for different AP-complexes) and it was stated in which AP comparisons these proteins were identified.

### GO term analysis

For GO Term analysis only proteins were used that were significantly enriched (p-value <0.05 and log_2_ fold change > 1) against both WT cells and cytosolic TurboID control (in case of KI vs overexpression, Fig. 2E) or against WT cell control and AP1μA/ AP2μ sample (for V5-TurboID-CLCa, Fig. 5). GO Term analysis was performed using the GO::TermFinder open source software^75^.

