## Supplementary figures and methods for "When less is more – A fast TurboID KI approach for high sensitivity endogenous interactome mapping"

**Supplementary information**

**Supplementary figures**

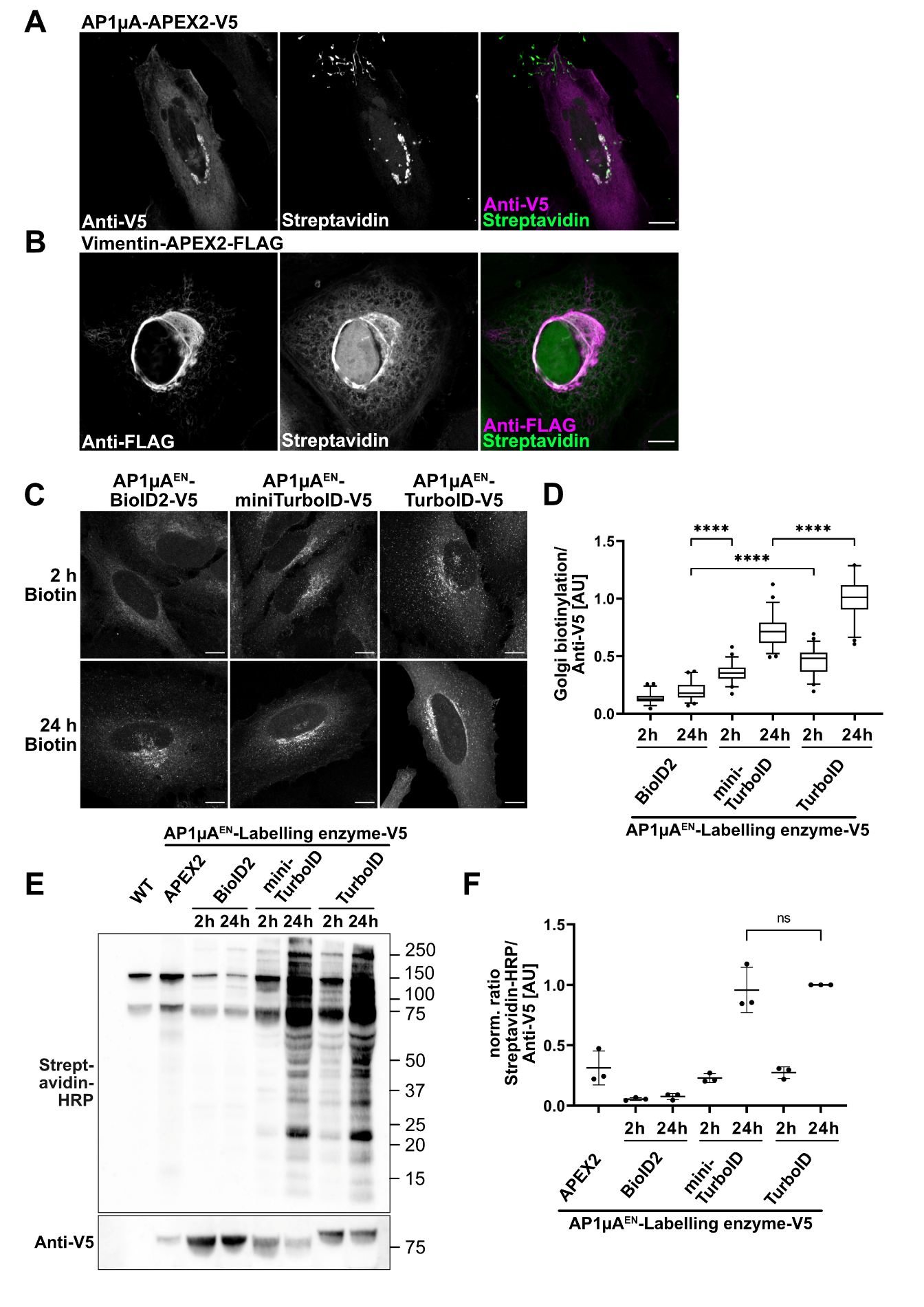

**Supplementary Fig. 1:** **Kinetics of labelling enzymes.**

**A:** Cell transiently overexpressing AP1µA-APEX2-V5 were fixed and stained with anti-V5 antibody to detect the fusion protein and streptavidin-AF488 to detect biotinylated proteins. Cells were incubated for 30 min with 500 µM biotin-phenol and labelling was induced for 1 min with H_2_O_2_. **B:** Cells transiently expressing Vimentin-APEX2-FLAG were fixed and stained with anti-FLAG antibody to detect the fusion proteins and streptavidin-AF488 to detect biotinylated proteins. Cells were treated as described in **A**. **C:** Detection of biotinylated proteins with streptavidin-AF488 in fixed cells expressing different biotin ligases. 50 μM biotin was added for either 2 h or 24 h before fixation. **D:** Ratio of biotinylated Golgi-localised proteins detected by streptavidin-AF488 to biotin ligase expression detected by anti-V5 antibody at the Golgi. Cells expressing the different biotin ligases endogenously fused to AP1μA were treated with 50 μM biotin for 2 h or 24 h and then fixed and prepared for microscopy as in **C**. For each condition 30-50 cells were analysed. All p-values from unpaired t-tests are <0.0001. **E:** Comparison of the labelling efficiency of different labelling enzymes endogenously fused to AP1μA on a western blot. Cells were treated with 50 μM biotin for 2 h or 24 h, WT (wild type) cells were treated for 24 h with 50 μM biotin and APEX2 were incubated for 30 min with 500 μM biotin-phenol and labelling was induced for 1 min with H_2_O_2_. Whole cell lysates were blotted with streptavidin-HRP to detect biotinylated proteins, and anti-V5 antibody to compare ligase expression. This experiment was performed three times with similar results. **F:** The ratio of biotinylated proteins (Streptavidin-HRP) to the expressed labelling enzyme (Anti-V5) was calculated for each line on the blot in **E** and then normalised to the TurboID value for each independent replicate. P-value is 0.71.

Scale bars are 10 μm.

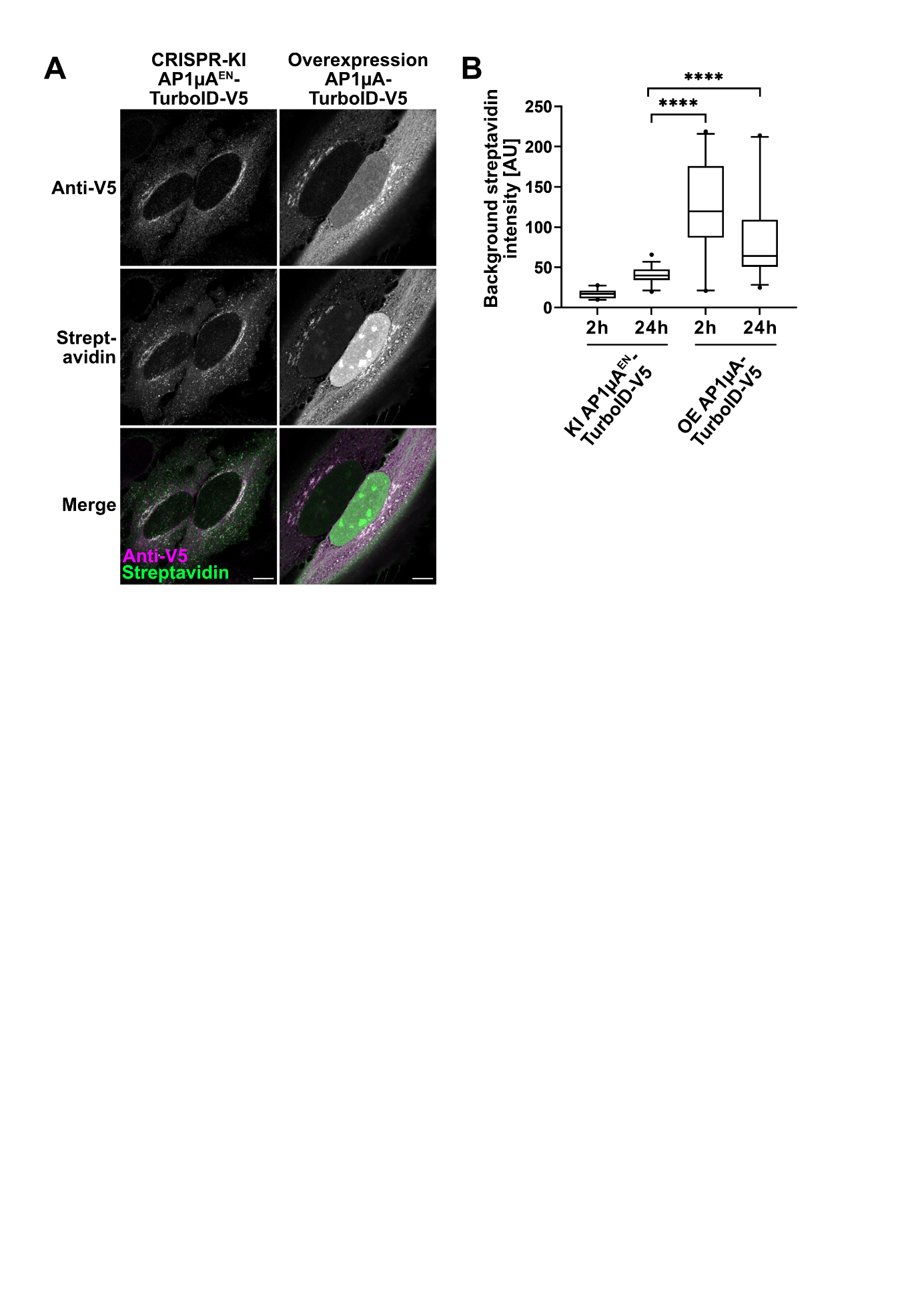

**Supplementary Fig. 2:** **Endogenous TurboID expression reduces unspecific labelling compared to transient overexpression.**

**A:** Comparison of cells either endogenously expressing AP1μA^EN^-TurboID-V5 or transiently overexpressing AP1μA-TurboID-V5. Cells were treated for 2 h with 50 μM biotin, fixed and stained with anti-V5 antibody to detect the labelling enzyme and streptavidin-AF488 to detect biotinylated proteins. **B:** Background biotinylation in cells either endogenously expressing AP1μA^EN^-TurboID-V5 (KI) or transiently overexpressing AP1μA-TurboID-V5 (OE). Background biotinylation was measured in cells that were treated with biotin for 2 h or 24 h by measuring streptavidin-AF488 signal intensity from a cytosolic area. At least 20 cells were analysed per condition. P-values are all <0.0001. Scale bars are 10 μm.

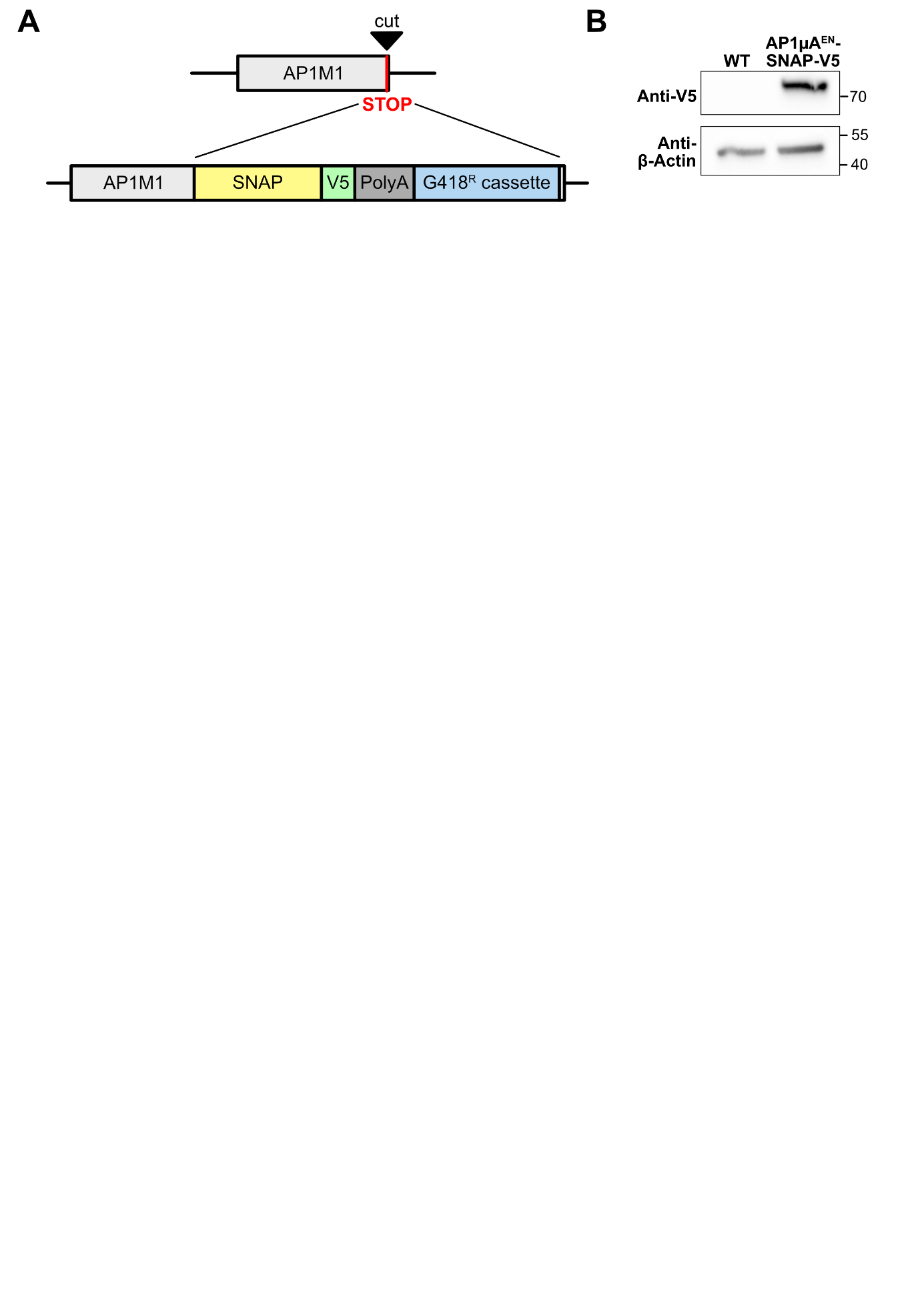

**Supplementary Fig. 3:** **Generation of an** **AP1µA^EN^-SNAP-V5 cell line.**

**A:** Scheme of KI strategy. AP1μA was C-terminally tagged with SNAP tag, a V5 tag and a resistance cassette that allows for rapid selection of positive cells. **B:** Blots of whole cell lysates to verify the AP1µA-SNAP-V5 KI by anti-V5 blotting.

**Supplementary methods**

*Generation of CRISPR-Cas9 knock-in cell lines*

All primers used in this section can be found in Supplementary Table 8.

The C-terminally tagged AP complex cell lines were generated following the strategy presented in Fig. 1A. The AP1M1 genomic locus (Gene ID 8907) was targeted shortly after the stop codon with the following guide RNA: 5´-CAGCCAACACCCCGGCCTCGGGG-3’ (PAM site underlined). The guide RNA was cloned into the SpCas9 pX459 plasmid (addgene plasmid #62988)^1^ by annealing oligos and ligation into the vector which was linearized with BbsI. The homology repair (HR) plasmid to generate the AP1µA-labelling enzyme cell lines contained ~1 Kb homology arms and was synthesized by Twist Bioscience. A glycin-serin linker (GSGSGSGSGS) and a BamHI and EcoRI site were designed between the two homology arms for the cloning of tags and resistance cassette as indicated in the schematic in Fig. 1A. The coding sequences of the different labelling enzymes were integrated between the homology arms, followed by a polyA sequence and a G418 resistance cassette that allows selection of positive edited cells with the drug G418/Geneticin (Thermo Fisher Scientific). The coding sequences of the various labelling enzymes were obtained via PCR from previously described vectors (addgene plasmids #66170, #74224, #107171, #107172)^2-4^ using sense primers with a BamHI restriction site and antisense primers with a NheI restriction site. The coding sequence for the SNAP-tag was obtained from pSNAPf vector (New England Biolabs) via PCR using a sense primer with a BamHI restriction site and an antisense primer with a NheI restriction site. The SV40 polyA sequence was amplified from pEGFP-C1 using a PolyA NheI sense and a PolyA NotI antisense primers. The G418 resistance cassette was amplified for pEGFP-C1 using a G418 NotI sense and G418 EcoRI antisense primers. The various fragments were cloned into the HR vector linearized with BamHI and EcoRI. Sequences of all primers used are provided in supplementary table 5. HeLa cells were transfected with 1 µg of pX459 plasmid with the AP1µA guide and 1 µg of HR-plasmid using FuGENE. G418 was added to the cells 3 days after transfection at a concentration of 1.5 mg/mL and media was exchanged every 2-3 days with new G418 at the same concentration until the selection was complete (after 7-10 days). After selection the cells were passaged every 2-3 days with a 0.5 mg/mL maintenance concentration of G418.

The AP2µ^EN^-TurboID-V5, AP3µA^EN^-TurboID-V5 and the AP4µ^EN^-TurboID-V5 cell lines were generated similar to the AP1µA^EN^-TurboID-V5 cell line.

The genomic locus of AP2M1 (Gene ID 1173) was targeted shortly after the stop coding with the guide RNA: 5´- ACTCGCTGCTAGCTGCCACTAGG -3’ (PAM site underlined). The guide RNA was cloned into the SpCas9 pX459 plasmid as described for AP1µA. The HR plasmid was synthetized by Twist Bioscience (~1 Kb homology arms). As for the AP1µA HR plasmid a glycin-serin linker and a BamHI and EcoRI site were designed between the two homology arms. To generate the AP2µ^EN^-TurboID-V5-PolyA-G418 HR plasmid the entire insert (TurboID-V5-PolyA-G418) was excised from the AP1µA^EN^-TurboID-V5-PolyA-G418 HR plasmid using the BamHI and the EcoRI site. The insert was cloned into the ordered AP2µ HR vector linearized with BamHI and EcoRI. The AP2µ^EN^-TurboID-V5-PolyA-G418 KI cell line was generated as described for AP1µA using the AP2µ guide vector and the AP2µ-TurboID-V5-PolyA-G418 HR plasmid.

AP3µA^EN^-TurboID-V5 and AP4µ^EN^-TurboID-V5 cell lines were generated as described for AP2µ^EN^-TurboID-V5. The genomic - locus of AP3M1 (Gene ID 26985) was targeted shortly after the stop coding with the guide RNA: 5´- TGGAAAACAAACTGGTCCTGAGG -3’ (PAM site underlined). The genomic locus of AP4M1 (Gene ID 9179) was targeted shortly after the stop coding with the guide RNA: 5´- GATCTGAGGCTCCCCAAACGAGG -3’ (PAM site underlined).

The AP4µ guide RNA was cloned into the SpCas9 pX330 plasmid (addgene plasmid #42230)^5^ by annealing oligos and ligation into the vector which was linearized with BbsI.

The N-terminal tagged CLCa-cells were generated following the strategy presented in Fig. 3A. The CLTA + genomic locus (Gene ID 1211) was targeted shortly after the start codon with the guide RNA: 5’- ATGGCTGAGCTGGATCCGTTCGG-3’. The guide RNA was cloned into the SpCas9 pX330 plasmid (addgene plasmid #42230)^5^ by annealing oligos and ligation into the vector which was linearized with BbsI. The CLCa-homology repair (HR) plasmid was synthesized by Twist Bioscience. It contained both homology arms (~1 Kb), a short N-terminal glycin-serin linker (SGSGSGSG) and a V5 coding sequence with a start codon. NheI and BamHI sites were designed between left homology arm and V5-sequence to integrate the resistance cassette and an EcoRI and SpeI site were designed for integration of the labelling enzymes. The coding sequences of the different labelling enzymes were obtained via PCR from previously described vectors (addgene plasmids #66170, #74224, #107171, #107172)^2-4^ using sense primers with an EcoRI restriction site and antisense primers with an SpeI restriction site. The G418 resistance cassette was amplified for pEGFP-C1 using a G418 NotI sense and G418 EcoRI antisense primers. A loxP sequence was integrated in both primers. The loxP-G418 fragment was cloned into the HR vector linearized with NheI and BamHI. In a second step the labelling enzyme fragments were cloned into the HR vector with the G418 cassette linearized with EcoRI and SpeI. HeLa cells were transfected with 1µg of pX330 plasmid with the CLCa-guide and 1 µg of HR-plasmid using FuGENE. After the selection with G418, the cells were again transfected with a CRE-transferase (addgene plasmid #11923)^6^ using the FuGENE transfection agent.

*Plasmid design of overexpression plasmids*

All primers used in this section can be found in Supplementary Table 8.

The sequences encoding for AP1µA-APEX2-V5, AP1µA-TurboID-V5 and cytosolic TurboID-V5 were all cloned into the pEGFP-N1 plasmid. The fragments of the labelling enzymes were obtained via PCR from the used HR plasmids using sense primers with a BamHI restriction site and antisense primers with a NotI restriction site. The eGFP was removed from the pEGFP-N1 vector by digestion with BamHI and NotI and the labelling enzyme fragments were inserted. The cDNA sequence of AP1µA including the same GS-linker that was used for the CRISPR-KI was synthesised by Twist Bioscience and the AP1µA-fragment including the linker was obtained via PCR using a sense primer with an EcoRI restriction site and an antisense primer with an BamHI restriction site. The AP1µA-fragment was cloned in the APEX2-V5 and TurboID-V5 vector linearized with EcoRI and BamHI.

The Vimentin-APEX2-FLAG plasmid (addgene #66170)^2^ was used for transient expression of Vimentin-APEX2.

The pHcgreen ITGB1-GFP plasmid (addgene #69804)^7^ was used for transient expression of ITGB1-eGFP.

The pEGFP VAMP7 (1-220) plasmid (addgene #42316)^8^ was used for transient expression of eGFP-VAMP7.

For transient expression of eGFP-SCYL2, the coding sequence of SCYL2 was obtained from pDONR223-SCYL2 (addgene #23458)^9^ via PCR using a sense primer with an EcoRI restriction site and an antisense primer with a BamHI restriction site. The SCYL2 fragment was cloned into the pEGFP-C1 vector using BamHI and EcoRI.

*Preparation of MS samples for the comparison of KI and overexpression*

The protocol for MS sample preparation is based on the protocol described in Cho et al.^10^. In total, three independent samples for each condition were prepared. For each KI-sample, AP1µA^EN^-TurboID-V5 cells were seeded into two T75 flasks (1.5 million cells per flask). For all other conditions (overexpression of AP1µA-TurboID-V5, cytosolic TurboID control and WT control), two 10 cm cell culture dishes were seeded with 1 million cells per dish. For transient overexpression of cytosolic TurboID or AP1µA-TurboID-V5, 18 h after seeding, cells were transfected with 4µg of either the AP1µA-TurboID-V5 pEGFP-N1 overexpression plasmid or the cytosolic TurboID-V5 pEGFP-N1 plasmid using FuGENE. 24 h after seeding, in all condition the medium was replaced with medium supplemented with 50 µM biotin. 48 h after seeding the samples were washed 5 times with ice-cold PBS and then detached in 4 mL PBS per flask/10 cm dish using a cell scraper and collected in a falcon tube. Cells were pelleted by centrifugation at 300g at 4°C for 3 min. The supernatant was removed and the pellet resuspended in 4 ml RIPA lysis buffer supplemented with 1x protease inhibitor and lysed for at least ten minutes on ice. The cell lysates were distributed into microcentrifuge tubes then clarified by centrifugation at 13000g at 4°C for 10 min. The clarified lysates were then mixed with 200 µl of streptavidin magnetic beads that were previously equilibrated twice with RIPA lysis buffer. The samples were distributed into fresh microcentrifuge tubes and incubated with the magnetic beads rotating at 4°C overnight. On the next day the beads were pooled and washed twice with RIPA lysis buffer (1 mL, 2 min), once with 1 M KCl (1 mL, 2 min) and then quickly once with 0.1 M Na_2_CO_3_ (1 mL, 10 s) and once with 2 M urea in 10 mM Tris-HCl (pH 8.0) (1 mL, 10 s). The beads were again washed twice in RIPA lysis buffer (1 mL, 2 min) and then transferred into fresh microcentrifuge tubes. They were then washed once with 50 mM Tris-HCl (pH 7.4) and twice with 2 M urea in 50 mM Tris-HCl (pH 7.4). The final wash buffer was then removed and the beads were resuspended in 80 µl of 2 M urea in 50 mM Tris-HCl (pH 7.4) with 1 mM DTT and 0.4 µg trypsin and incubated for 1 h shaking at 1000 r.p.m.. The supernatant was transferred into fresh microcentrifuge tubes and the beads were washed twice with 60 µl of 2 M urea in 50 mM Tris-HCl (pH 7.4). The washes were combined with the supernatant, DTT was added to a final concentration of 4 mM and the samples were incubated for 30 min at 25°C shaking at 1000 r.p.m.. Next, iodoacetamide was added to final concentration of 10 mM and the samples were incubated in the dark at 25°C for 45 min while shaking at 1000 r.p.m.. Finally, another 0.5 µg of trypsin were added and the digestion was proceeded overnight at 25°C and shaking at 700 r.p.m.. On the next day, the digestion was stopped by acidification with formic acid to a final concentration of 1% (vol/vol). The peptides were prepared for MS using SDB-stage tips.

*Preparation of MS samples for comparison of different AP complexes*

For each sample (4 independent samples in total), cells of the different KI cell lines (AP1µA^EN^-TurboID-V5, AP2µ^EN^-TurboID-V5, AP3µA^EN^-TurboID-V5 and AP4µA^EN^-TurboID-V5) were seeded into two 15 cm cell culture dishes (1.5 million cells per dish). Due to the low abundance of AP-4 a third 15 cm cell culture dish with 1.5 million cells was used. As a control HeLa WT cells were seeded into two 15 cm cell culture dishes (1.5 million cells per dish). 24 h after seeding, the medium was replaced with medium supplemented with 50 µM biotin for another 24 h. For harvesting and enrichment the protocol was comparable to the one described above. To ensure sufficient cell lysis and full disruption of membranes, all pellets were resuspended in 4 mL RIPA lysis buffer supplemented with 1x protease inhibitor and lysed for 30 minutes on ice. During this time, the cell membranes were disrupted by passing the cells 10 times (5 strokes) through a 24 G needle.

*Preparation of MS samples for V5-TurboID-CLCa*

For each sample (4 independent samples in total), V5-TurboID-CLCa^EN^ cells were seeded into two 15 cm cell culture dishes (1.5 million cells per dish). As a negative control HeLa WT cells were seeded in two 15 cm cell culture dishes (1.5 million cells per dish). 24 h after seeding, in all condition the medium was replaced with medium supplemented with 50 µM biotin for another 24 h. For harvesting and enrichment the protocol was comparable to the one described above.

*Sample clean-up by SDB-STAGE tips*

Salts and detergents were removed before LC-MS analysis by SDB-RPS StageTips (Empore™ 2241). Stage tips were prepared as described previously ^11^. For clean-up, condition StageTips with 200 μL of 100% MeOH, 200 μL of solvent 1 (80% Acetonitrile, 0.1% formic acid), and twice with 200 μL of solvent 2 (0.1% formic acid). Load acidified peptides onto the conditioned StageTips and wash twice with 200 μL of solvent 2 and once with 200 µL solvent 1. Elute the peptides from the StageTips with 20 μL of elution buffer (5% NH_4_OH in 60% acetonitrile) and vacuum centrifuge the samples until completely dry.

*Nano liquid chromatography-mass spectrometry (LC-MS) and data analysis*

Peptides were reconstituted in 10 µl of 0.05% trifluoroacetic acid (TFA), 4% acetonitrile, and 5 µl were analyzed by an Ultimate 3000 reversed-phase capillary nano liquid chromatography system connected to a Q Exactive HF mass spectrometer (Thermo Fisher Scientific). Samples were injected and concentrated on a trap column (PepMap100 C18, 3 µm, 100 Å, 75 µm i.d. x 2 cm, Thermo Fisher Scientific) equilibrated with 0.05% TFA in water. After switching the trap column inline, LC separations were performed on a capillary column (Acclaim PepMap100 C18, 2 µm, 100 Å, 75 µm i.d. x 25 cm, Thermo Fisher Scientific) at an eluent flow rate of 300 nl/min. Mobile phase A contained 0.1 % formic acid in water, and mobile phase B contained 0.1% formic acid in 80 % acetonitrile / 20% water. The column was pre-equilibrated with 5% mobile phase B followed by an increase of 5–44% mobile phase B in 100 min. Mass spectra were acquired in a data-dependent mode utilising a single MS survey scan (m/z 300–1650) with a resolution of 60,000, and MS/MS scans of the 15 most intense precursor ions with a resolution of 15,000. The dynamic exclusion time was set to 20 seconds and automatic gain control was set to 3x10^6^ and 1x10^5^ for MS and MS/MS scans, respectively.

MS and MS/MS raw data were analysed using the MaxQuant software package (version 2.0.2.0) with implemented Andromeda peptide search engine^12^. Data were searched against the human reference proteome downloaded from Uniprot (79,684 proteins, taxonomy 9606, last modified April 18, 2022) using the default parameters except for enabling the options label-free *quantification (LFQ) and match between runs.* Filtering and statistical analysis was carried out using the software Perseus version 1.6.14 ^13^.

Only proteins which were identified with LFQ intensity values in all 3 replicates within at least one experimental group (for AP1µA-TurboID KI vs overexpression, Fig. 2) or in at least 3 out of 4 replicates (CLCa MS and AP complex interactome mapping, Fig 3, 5), were used for downstream analysis. Missing values were replaced from normal distribution (imputation) using the default settings in Perseus (width 0.3, down shift 1.8). Mean log_2_ fold protein LFQ intensity differences between experimental groups calculated in Perseus using student’s t-tests with permutation-based FDR of 0.05 to generate the adjusted p-values (q-values).

*Volcano plots*

Volcano plots were created by plotting the -log_10_ p-value against the mean log_2_ fold protein LFQ intensity differences. The volcano plots in Fig. 2 show all proteins that were significantly enriched over the WT control. For the volcano plots in Fig. 3 and 5 all proteins were included in the volcano plot representation to prevent exclusion of potential interactors for one of the two bait proteins that are compared.

**Supplementary tables**

**Supplementary Table 7: Antibody and Streptavidin conjugates**

| Antibody/Conjugate | Supplier | Used concentration |
| --- | --- | --- |
| V5-Tag (D3H8Q) Rabbit | Cell Signaling | 1:1000 for IF and WB |
| DYKDDDDK Tag (D6W5B) Rabbit | Cell Signaling | 1:1000 for IF |
| β-Actin (8H10D10) Mouse | Cell Signaling | 1:1000 for WB |
| Anti-AP1M1 (ab230273) Rabbit | Abcam | 1:1000 for WB |
| Anti-CHC Mouse | Novus Biologicals | 1:1000 for IF |
| Anti-p230 Mouse | BD transduction laboratories | 1:1000 for IF |
| Anti-Rabbit- Atto647N Goat | Sigma-Aldrich | 1:1000 for IF |
| Anti-Mouse- Alexa594 Goat | Invitrogen | 1:1000 for IF |
| Anti-Rabbit- HRP Goat (ab6721) | Abcam | 1:5000 for WB |
| Anti-Mouse- HRP Goat (ab6789) | Abcam | 1:5000 for WB |
| Anti-Rabbit- HRP Goat (ab6721) | Abcam | 1:5000 for WB |
| Streptavidin, HRP conjugate (N100) | Thermo Scientific | 1:4000 for WB |
| Streptavidin, Alexa Fluor™ 488 conjugate (S11223) | Invitrogen | 1:1000 for IF |
| Streptavidin, STARORANGE conjugate | Abberior | 1:500 for IF |

**Supplementary Table 8: Primers**

| Primer Name | Primer Sequence (5’-3’) |
| --- | --- |
| PolyA sequence: |  |
| PolyA NheI sense | AGTTCGCTAGCCCGCGACTCTAGATCATAATCAGC |
| PolyA NotI antisense | TGCACGCGGCCGCCTTACAATTTACGCCTTAAGATACA |
| G418 cassette for C-terminal KI: |  |
| G418 NotI sense | CGTCGGCGGCCGCCCTGAGGCGGAAAGAACCAGCTGTGGAATGTGTGTCAGTTAG |
| G418 EcoRI antisense | TCATGGAATTCTTTATTCTGTCTTTTTATTGCCGTC |
| Labelling enzymes for C-terminal KI: |  |
| APEX2 BamHI sense | GATGGGGATCCATGGGGAAATCATACCCAACAGTGTCCG |
| APEX2 V5 NheI antisense | CTAACGCTAGCTCAGGTGCTGTCCAGGCCCAGCAGGGGGTTGGGGATGGGCTTGCCGGCGTCGGCAAATCCCAGTTCTGAC |
| BioID2 BamHI sense | GATGGGGATCCATGTTCAAGAACCTGATCTGGCTGAAGG |
| BioID2 V5 NheI antisense | CTAACGCTAGCTCAGGTGCTGTCCAGGCCCAGCAGGGGGTTGGGGATGGGCTTGCCGCTTCTTCTCAGGCTGAACTCGCCG |
| TurboID BamHI sense | GATGGGGATCCATGAAAGACAATACTGTGCCTCTGAAGC |
| MiniTurboID BamHI sense | GATGGGGATCCATGATCCCGCTGCTGAACGCTAAACAGA |
| TurboID/MiniTurboID V5 NheI antisense | CTAACGCTAGCTCAGGTGCTGTCCAGGCCCAGCAGGGGGTTGGGGATGGGCTTGCCCTTTTCGGCAGACCGCAGACTGATT |
| SNAP primer for C-terminal KI: |  |
| SNAP BamHI sense | ctgatGGATCCGACAAAGACTGCGAAATGAAGCGCA |
| SNAP V5 NheI antisense | TACAAGCTAGCTCAGGTGCTGTCCAGGCCCAGCAGGGGGTTGGGGATGGGCTTGCCACCCAGCCCAGGCTTGCCCAGTCTG |
| G418 cassette with loxP sites for N-terminal KI: |  |
| G418 loxP NheI sense | ATGTCGCTAGCATAACTTCGTATAGCATACATTATACGAAGTTATCCTGAGGCGGAAAGAACCAGCTGTGGAATGTGTGTCAGTTAG |
| G418 loxP BamHI antisense | TGCACGGATCCGTGATAACTTCGTATAATGTATGCTATACGAAGTTATTTTATTCTGTCTTTTTATTGCCGTCATAGCGCGGGTT |
| Labelling enzymes for N-terminal KI: |  |
| APEX2 EcoRI sense | CTAACGAATTCGGGAAATCATACCCAACAGTGTCCG |
| APEX2 SpeI antisense | CTAACACTAGTGGCGTCGGCAAATCCCAGTTCTGAC |
| BioID2 EcoRI sense | CTAACGAATTCTTCAAGAACCTGATCTGGCTGAAGG |
| BioID2 SpeI antisense | CTAACACTAGTGCTTCTTCTCAGGCTGAACTCGCCG |
| TurboID EcoRI sense | CTAACGAATTCAAAGACAATACTGTGCCTCTGAAGC |
| MiniTurboID EcoRI sense | CTAACGAATTCATCCCGCTGCTGAACGCTAAACAGA |
| TurboID/ MiniTurboID SpeI antisense | GTCACACTAGTCTTTTCGGCAGACCGCAGACTGATT |
| Overexpression plasmids: |  |
| APEX2 BamHI sense | CTAACGGATCCGGGAAATCATACCCAACAGTGTCCG |
| (APEX2) V5 NotI antisense | CTAACGCGGCCGCTCAGGTGCTGTCCAGGCCCAGCAGGGG |
| TurboID BamHI sense | AGTCAGGATCCATGAAAGACAATACTGTGCCTCTGAAGCTG |
| TurboID V5 NotI antisense | TAGCTGCGGCCGCTCAGGTGCTGTCCAGGCCCAGCAGGGGGTTGGGGATGGGCTTG |
| AP1µA EcoRI sense | GATGGGAATTCATGTCCGCCAGCGCCGTCT |
| AP1µA linker BamHI antisense | TGTGCGGATCCTGAGCCGGAACCAGAGCCTGACCCT |
| SCYL2 EcoRI sense | TACCGGAATTCTatggagtccatgcttaataA |
| SCYL2 BamHI antisense | TACCGGGATCCtcacccaaaaagatcttttaaat |
